# I^2^SIM: Boosting High-Fidelity Isotropic Super-Resolution with Image Interference and Spatial-Spectral Optimization

**DOI:** 10.1101/2024.12.18.629092

**Authors:** Enxing He, Yile Sun, Hongfei Zhu, Xinxun Yang, Lu Yin, Yubing Han, Cuifang Kuang, Xu Liu

## Abstract

Spatial resolution is crucial for imaging subcellular structures. The advent of three-dimensional structured illumination microscopy (3D-SIM) greatly benefits the biology community, providing a powerful tool for imaging organelles with a two-fold resolution enhancement in all three dimensions. However, the axial resolution of 3D-SIM is limited to around 300 nm, which is inferior to its lateral resolution. Here, a novel method called image interference SIM (I^2^SIM) is reported, which utilizes two oppositely positioned objectives to detect fluorescence emission interference under three-beam excitation. By incorporating spectral modulation and spatial domain Frobenius-Hessian optimization, I^2^SIM achieves an axial resolution approximately twice that of 3D-SIM, reaching around 130 nm. Furthermore, the potential of I^2^SIM for imaging subcellular structures is demonstrated on various biological samples, including microtubules, actin filaments, and mitochondrial outer membranes. The enhanced optical sectioning capability can be utilized to resolve axial structures that are challenging to discern using ordinary 3D-SIM.

## 1 Introduction

Observation of subcellular structures at high spatial resolution plays a crucial role in cell biology. Due to the existence of diffraction limit, it is challenging to clearly discern fine structures in three dimensions. The advent of super-resolution techniques breaks down this barrier and offers new perspectives for studying the behavior of organelles. Among these, structured illumination microscopy (SIM) is widely utilized because of its advantages of low phototoxicity and high temporal resolution. By exciting the sample with two symmetrically inclined plane waves, SIM can encode the high-frequency information into the diffraction-limited passband, and doubles the resolution (120 nm in lateral) by demodulating and stitching together low- and high-frequency components. [1, 2, 3, 4]. Compared to its 2D counterpart, 3D-SIM introduces an additional central illumination beam, which generates variations in the axial illumination field, enabling a two-fold axial super-resolution of approximately 300 nm. The performance of SIM is closely tied to that of the reconstruction algorithms. Significant efforts have been devoted to the development of these algorithms [5, 6, 7, 8, 9, 10, 11, 12, 13], enhancing image fidelity and improving optical sectioning effect. The synchronized development of hardware and software has enabled 3D-SIM to reveal a multitude of biological mechanisms [14, 15, 16, 17].

However, a major limitation of 3D-SIM is its anisotropic resolution, with the axial resolution being substantially lower than the lateral. Achieving isotropic resolution of 3D-SIM has naturally become an important area of focus. Indeed, the resolution of SIM is directly linked to the highest spatial frequency component. Therefore, improving axial resolution is equivalent to increasing the maximum detectable axial frequency. This objective can be accomplished through illumination modulation, detection modulation, or a combination of both. From the perspective of increasing illumination frequency, four-beam SIM enhances axial resolution to approximately 160 nm by positioning a mirror against the objective lens to reflect the central illumination beam, thereby generating a four-beam illumination in 3D-SIM configuration [18]. The integration of illumination modulation and detection modulation pushes the axial resolution even further. I^5^S employed six excitation beams for illumination and collected bidirectional fluorescence using two opposite objectives to extend the supporting region of the optical transfer function (OTF) [19, 20, 21]. Nevertheless, enhancing resolution inevitably results in increased system complexity. Due to the intricate structured illumination fields and complicated frequency shifts, the practical application of I^5^S remains challenging. Although four-beam SIM simplified the acquisition setup using wide-field detection, it still needs fine-tuning at the illumination end. And a common challenge in reconstruction is the presence of missing or low-value components in the synthesized spectrum. This unevenness leads to artifacts and noise in the reconstructed images, ultimately degrading image quality.

Here we propose a novel method, I^2^SIM, which utilizes opposite objectives based on I^2^M [22, 23, 24] and applies three excitation beams to avoid cumbersome adjustments at the illumination end. From the perspective of enhancing reconstruction algorithm performance, inspired by Hessian-SIM [25, 26] and HiFi-SIM, joint spectral modulation and spatial optimization are further employed to improve the imaging quality, eliminating artifacts and suppressing noises. First, spectral filters are constructed to uniformize the synthesized spectrum and minimize side lobes. Second, spatial domain Frobenius Hessian optimization is applied to the SIM result with the uniformly synthesized spectrum as a fidelity term to further reduce noise and artifacts, enhancing the continuity of the imaging outcomes. Our approach extends axial resolution to approximately 130 nm under 560 nm excitation, while maintaining a lateral resolution of around 100 nm and providing extraordinary image quality. We tested our method in microtubules, actin filaments, and mitochondrial outer membranes.

## 2 Results

### 2.1 Principle

#### 2.1.1 Extension of Optical Transfer Function

Dual-objective detection, also known as 4Pi configuration, naturally enhances the axial resolution in wide-field imaging due to the axial extension of OTF. Through interference of forward and backward fluorescence, the supporting region of OTF in the *k*_*z*_ axis can be extended from [1− *cos*(*sin*^*−*1^(NA/*n*))] · *nk*_*em*_ to 2*nk*_*em*_, where NA denotes the numerical aperture, *n* is the refractive index of the immersion medium, and *k*_*em*_ is the wave vector of fluorescence in vacuum. I^2^SIM adopts the three-beam excitation of 3D-SIM (**Figure 1a**) to extend the 4Pi OTF in both lateral and axial directions within Fourier domain, while also compensating for the missing frequency components inherent in the 4Pi OTF (**Figure 1a**). Spatial-spectral optimization balances the relative intensities of the high and low-frequency components of the synthesized spectrum, resulting in a smoother extended OTF that enables isotropically enhanced resolution and reduces artifacts. A comparison of the synthesized OTFs of 3D-SIM and I^2^SIM can be found in **Figure 1c**, where I^2^SIM improves the axial resolution by a factor of two compared to 3D-SIM.

**Figure 1:**
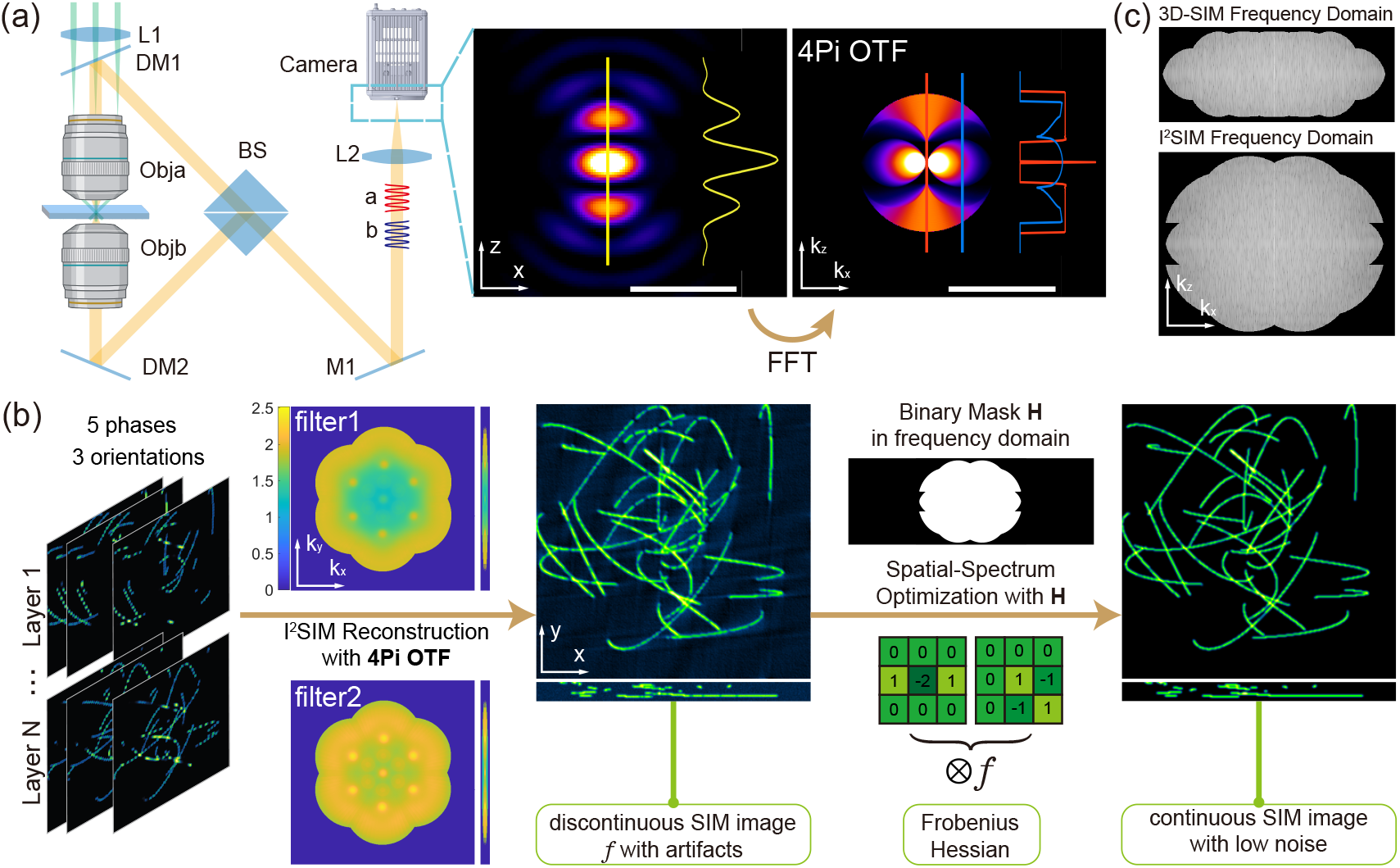
Principle of I^2^SIM. a) (Left)Structure schematic diagram of 4Pi-SIM. The sample is illuminated by three beams the same as 3D-SIM through one objective and fluorescence detected by both objectives will interfere with each other on CMOS; (Middle)4Pi PSF on CMOS; (Right)The corresponding OTF. The line profile of PSF is normalized and Profiles of OTF were calculated by normalized *log*(*abs*(*Intensity*) + 1). b) Algorithm flow. c) Comparison of OTF supporting region between 3D-SIM and I^2^SIM.

#### 2.1.2 Spectral Modulation

In ordinary SIM reconstruction, the synthesized spectrum method based on Wiener filtering (Wiener-SIM) is utilized. The spectrum is the recombination of different frequency components in the Fourier domain, but exhibiting an uneven intensity profile. The peaks and kinks of the synthetic OTF in Wiener-SIM will lead to side lobes in the space domain, reducing resolution and increasing artifacts. Inspired from Open-3DSIM [27], we applied two spectrum filters to reshape the equivalent OTF and adjustedmit to suit for I^2^SIM. The cooperation of filters makes the 3D spectrum more ideal, smoother, and more uniform, addressing the issues mentioned above. The pipeline is shown in Figure 1b. Spectrum optimization is specifically divided into two steps. First, after the raw image stack is captured, a notch filter is applied to the spectrum separated by SIM reconstruction to suppress the high-frequency peaks and minimize background noise. Then we obtained the preliminarily processed and synthesized spectrum 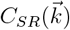. More detailed derivations of 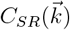 can be found in Note S2 of the Supporting Information. Second, OTF_*notch*_ is built to design the filters for spectrum optimization to improve the patchy features in *C*_*SR*_(*k*).

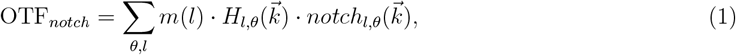

where notch_*l,θ*_ 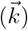 and *H*_*l,θ*_ 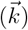 are the notch-filter and leaky-cone-shaped optical transfer function shifted to the *l*-th place in the frequency domain, respectively. *θ* denotes the pattern orientation in the lateral plane. *l* is the symbol of the order of the separated frequency components. The mathematical model of them can be found in Note S2 of the Supporting Information. *m*(*l*) is the weight coefficient of different Fourier orders. Then, OTF_*notch*_ is used to build frequency domain *filter*_1_ and *filter*_2_. Specifically,

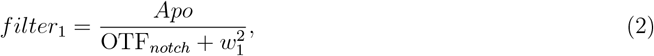

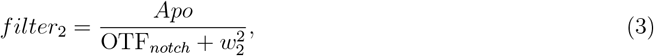

where *Apo* is the apodization function in the 3D frequency domain. *w*_1_ and *w*_2_ are the parameters to design filters, respectively. Applying *filter*_1_ can suppress the patchy features and high-frequency noise, but decrease weak information, especially high-frequency information, at the same time. With a relatively smaller parameter *w*_2_, *filter*_2_ can increase the proportion of high-frequency components, thus maintain the weak information.

The filters are applied to the directly combined SIM spectrum *C*_*SR*_ obtained in the first step to suppress spectral side lobes, thereby reducing artifacts and enhancing the quality of reconstruction.

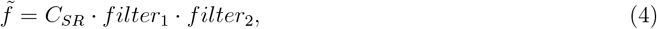

where 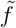 is the final SIM spectrum output of open-3DSIM with less artifacts.

#### 2.1.3 Spatial Optimization

Although filters can increase resolution and surpass noise and side lobes, artifacts and noises still exist due to the unsmoothed synthetic OTF. To solve this problem, space-domain optimization using the Frobenius norm of Hessian and l_1_ norm as regularization is applied. Upon finishing SIM reconstruction, we can construct the following loss function:

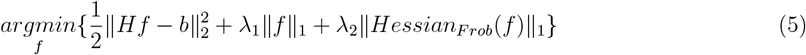

Here, *f* is the inverse Fourier Transform of 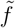. The operator *H* is a binary frequency mask which shares the same region as reconstructed effective OTF to make sure the fidelity is considered only in physically detected area. ∥ · ∥_1_ and ∥ · ∥_2_ are *l*_1_ and *l*_2_ norms, respectively. *l*_1_ norm guarantees the sparsity andm Frobenius norm of Hessian recovers the continuity of sample with better SNR increase (ISNR) [28], respectively. The Frobenius norm of Hessian can be represented as:

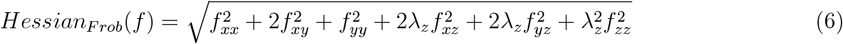

Here *f*_*xx*_ = ∂^2^*f*/∂*x*^2^, and similar operations are performed for others. λ_1_, λ_2_, and λ_*z*_ adjust the weights of various terms, balancing fidelity and regularization. By optimizing the loss function using the Alternating Direction Method of Multipliers (ADMM) [29] iteratively which is further discussed in Supporting Information Note S1, the recovered OTF will have a more uniformly extended supporting region than ordinary 3D-SIM as shown in Figure 1c, ensuring ideal reconstruction quality.

### 2.2 Simulation and Calibration

Simulated three-dimensional fluorspheres and microtubules were used to verify the effectiveness of I^2^SIM. Fluorspheres were generated as ideal points and convolved by wide-field PSF and 4Pi PSF with 0 phase difference, respectively. Both PSFs were generated using vectorial diffraction theory with a 1.45 NA objective and a 1.518 refractive index. The axial range was from -750 nm to 750 nm with 31 layers. As shown in **Figure 2a**, I^2^SIM shared the same lateral resolution as 3D-SIM. The axial resolution of wide-field, 3D-SIM and I^2^SIM were compared on the right side of Figure 2a, the axial performance of fluorspheres were dramatically improved and OTF supporting area’s max value was extended from ∼ 1.4*k*_*em*_ to ∼ 2.7*k*_*em*_. The full-width at half-maximum (FWHM) values for wide-field, 3D-SIM, and I^2^SIM are 497.1 nm, 284.4 nm, and 133.0 nm under center emission wavelength of 590 nm, respectively. Simulated microtubules were also reconstructed to test the performance under the same imaging parameters with 21 layers collected in axial direction. As shown in Figure 2c, the spatial resolution of I^2^SIM is closer to isotropic and can easily distinguish the axial multi-layers of simulated microtubules. It should also be emphasizes that, although 3D-SIM and I^2^SIM share the same lateral resolution, our approach exhibits fewer artifacts in each slice. This improvement is attributed to the higher axial resolution, which mitigates the influence of adjacent slices, thus showing clearer axial structures (Figure 2c).

**Figure 2:**
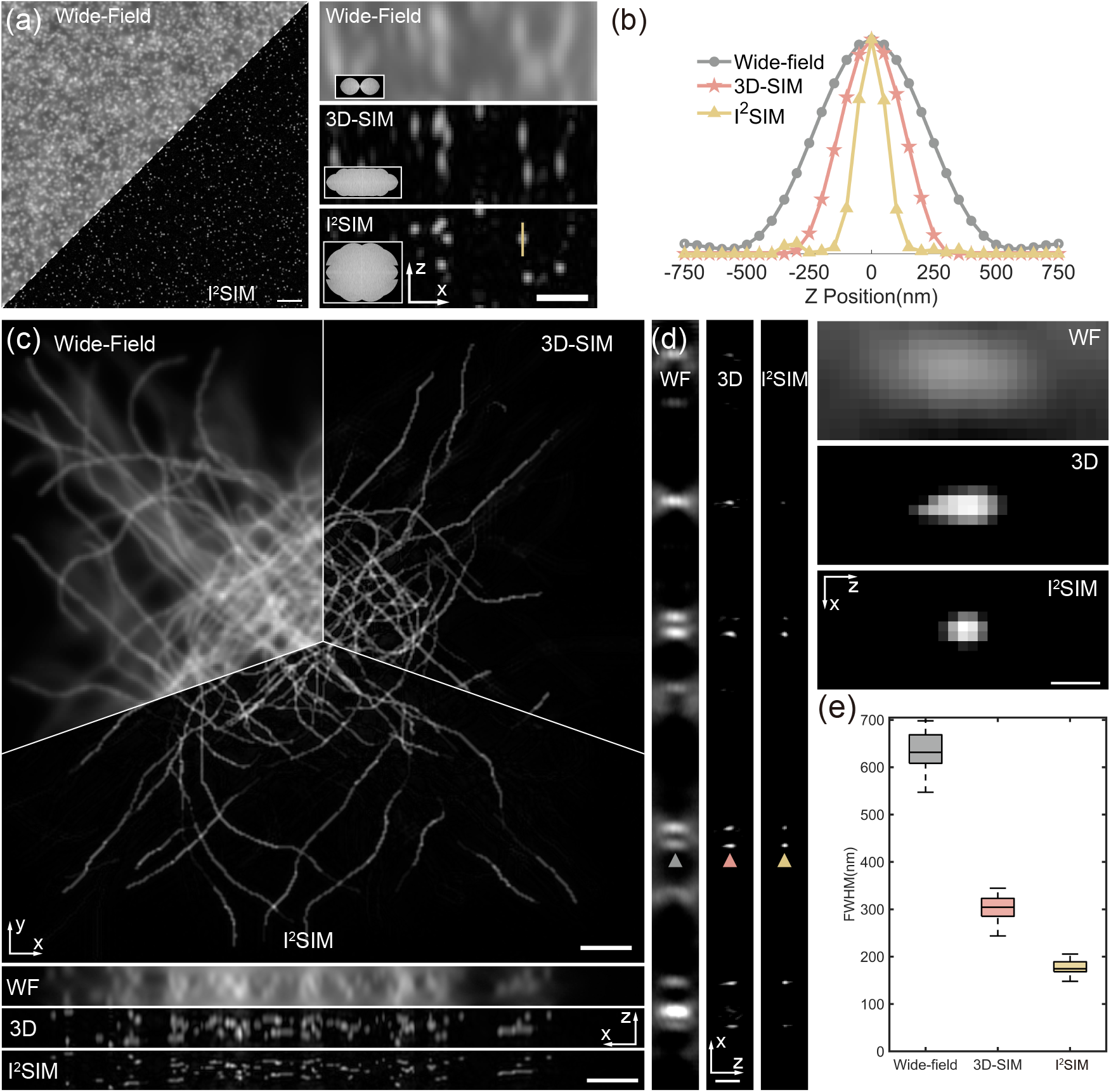
Simulation and beads experiment. a) (Left)One layer of simulated beads reconstruction result; (Right)Axial cross-sectional views of simulated beads. Insets show the magnitude of OTFs(*k*_*x*_*/k*_*z*_plane) derived from images. b) Line profiles corresponding to bead images shown on the right side of a), taken along the vertical line in a). c) Reconstruction results of simulated microtubules, including the comparison of the wide-field, open3D-SIM, and our method. d) (Left)One layer of reconstruction results of 100 nm beads in x-z plane; (Right)Higher magnification views of bead highlighted by colored arrowheads in Left. e) Quantification of axial FWHM for n = 76, 100 and 100 beads for wide-field microscopy, 3D-SIM and I^2^SIM, respectively.Whiskers: maximum and minimum; center lines: medians; bounds of box: 75th and 25th percentiles. Scale bars, 2 *µ*m in (a)(Left), 800 nm in (a)(Right), 2 *µ*m in (c), 800 nm in (d)(Left), 250 nm in (d)(Right).

Furthermore, We also examined reconstruction performance on a high-density Fluorspheres (FluoSpheres beads, F8801) with 100 nm diameter under 560 nm excitation wavelength. In Figure 2d, as expected, I^2^SIM maintained the ∼ 2-fold lateral resolution enhancement of 3D-SIM while offering ∼ 2-fold-better axial resolution than 3D-SIM. It can be observed that the average FWHM of wide-field, 3D-SIM, and I^2^SIM were 633.7 nm, 302.3 nm, and 178.4 nm respectively. Taking into account the size of the fluorspheres, this result is in line with our expectations.

### 2.3 Cell Imaging

To verify the performance of I^2^SIM in applications, several subcellular structures with more complicated imaging conditions were observed. Fixed microtubule networks of BSC-1 cells labeled with Abberior STAR red were first tested (**Figure 3**). The spatial illumination pattern was generated by a 647 nm Laser and centered at the focal plane of both objective lenses. for each layer, 15 images(5 phases per orientation × 3 orientation) were collected and then moved the z-piezo stage every 50 nm interval to achieve axial scanning and focus on another layer. Depth-encoded images revealed that microtubules were primarily concentrated in the upper layer of the field of view. Furthermore, the enlarged region confirmed that I^2^SIM matched the lateral resolution of 3D-SIM and experienced reduced axial interference (Figure 3a). In axial view, wide-field microscopy could barely discern the closely spaced microtubules, the intervals of which were more distinctly visible in 3D-SIM. Further, by employing I^2^SIM, structures with a spacing of 200 nm in axial direction can be clearly resolved, manifesting the ability of unveiling axial detail otherwise blurred by diffraction (Figure 3b, c). Similar sample with fibrous structure (for example, actin filaments) can be found in **Figure S1**.

**Figure 3:**
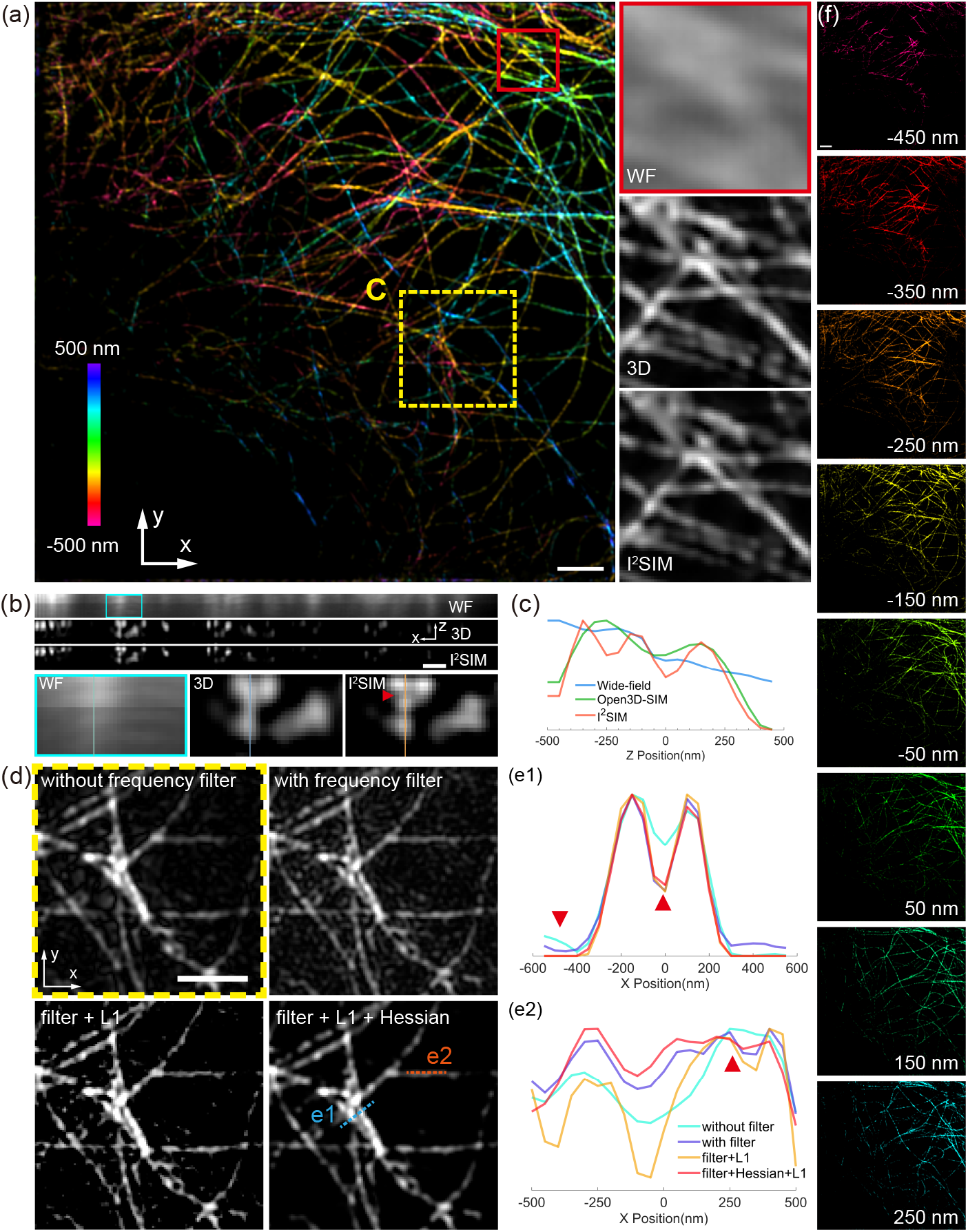
I^2^SIM enables near-isotropic imaging of microtubules. a) (Left)Maximum intensity projection of fixed BSC-1 cells labeled with the Abberior STAR Red. Image is depth coded as indicated; (Right)Higher magnification lateral views(single planes) corresponding to the red rectangle in (Left) are shown, comparing wide-field microscopy(upper), 3D-SIM(middle), and 4Pi-SIM(lower). b) (Top)Axial cross-sectional views of microtubules in a); (Middle)Higher magnification views corresponding to the rectangle indicated in the wide-field (WF) image at the top. c) Line profiles corresponding to the vertical line in b). d) Higher magnification lateral views corresponding to the yellow dashed rectangle in a), comparing unmodified image, frequency-filtered image, image modified by frequency filter and sparsity regularization, and image modified by frequency filter and sparsity plus Frobenius Hessian regularization. e) Line profiles taken along dashed lines e1 and e2 in d). f) Continuous x-y section of the 3D volume from -450 to 250 nm depth with a 100-nm axial interval, illustrating the changing morphology of microtubule networks. Scale bars, 2 *µ*m (a, f); 1 *µ*m (b); 1.5 *µ*m (d).

The effectiveness of the algorithm was also tested. By employing frequency domain filters in the open 3D-SIM reconstruction process, side lobes were surpassed, resulting in structures with clearer edges and enhanced resolution. *l*_1_ norm was used to further mitigate noise and improve the resolution, but it can lead to discontinuities in the structure. The purpose of the Frobenius Hessian norm was to correct these discontinuities and maintain spatial resolution, achieving an improvement in image quality (Figure 3d). Profiles in (Figure 3e1) show the suppression of noise and the preservation of resolution, while profiles along the direction of the structure (Figure 3e2) show a smoothing of intensity variations and an enhancement of continuity. By demonstrating several slices in layer stack (Figure 3f), the orientation and spatial distribution of microtubule growth can be observed. The nearer to the cell center, the microtubule distribution was closer to the upper layer, showing an upward trend.

To test the ability of I^2^SIM in the more complicated architecture, we imaged immunolabeled outer membranes of mitochondria in fixed U2OS cells (**Figure 4a**). Due to the diffraction limit, wide-field microscopy can hardly discern the hollow structure of the membranes while 3D-SIM and I^2^SIM can reveal clearly (Figure 4b). The axial resolution enhancement enabled I^2^SIM to reconstruct more intricate membrane invaginations that appeared indistinct or badly blurred in 3D-SIM (Figure 4b, c). Through precise axial scanning, I^2^SIM can build up the three-dimensional distribution and morphology of the mitochondrial outer membrane with nearly isotropic resolution (Figure 4d) and enabled visualization of fine mitochondrial substructure in both axial (Figure 4e) and lateral views.

**Figure 4:**
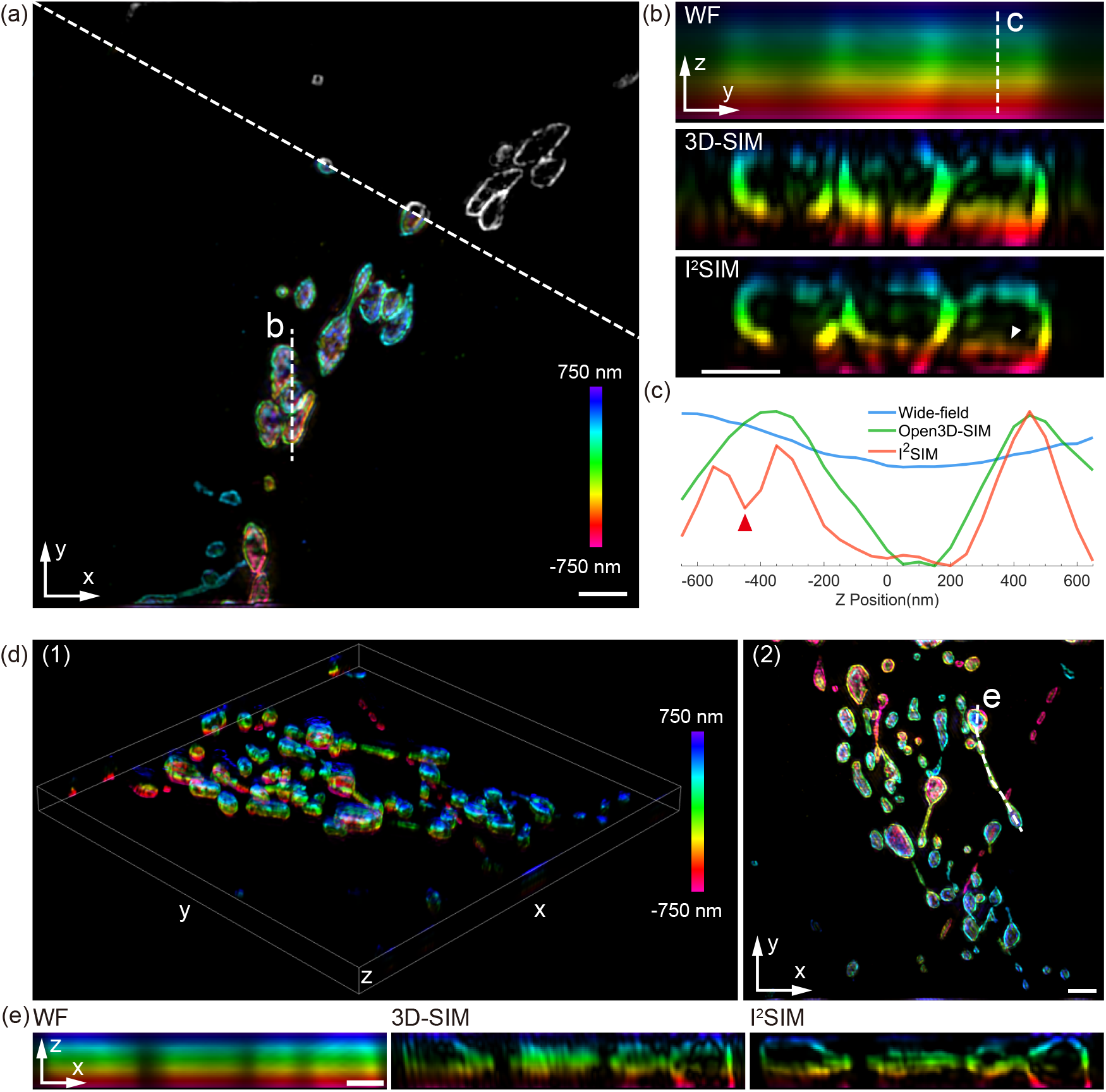
I^2^SIM revealed 3D architecture of the mitochondrial outer membrane. a) I^2^SIM image (maximum intensity projection) of fixed U2OS cells immunolabeled with Alexa Fluor 546. The right upper shows the x-y cross-section at z = 250 nm. b) Axial views corresponding to the white dashed line in (a), comparing wide-field microscopy (WF), 3D-SIM, I^2^SIM. c) Line profiles corresponding to the white dashed line in (b) for WF, 3D-SIM, and I^2^SIM. d) (Left)A 3D rendering of mitochondrial outer membrane; (Right)Maximum intensity projection in x-y view. e) Axial views corresponding to the white dashed line in (d2), comparing wide-field microscopy (WF), 3D-SIM, I^2^SIM. Scale bars, 2 *µ*m (a, d2); 1 *µ*m (b, e).

## 3 Discussion

In order to achieve high-quality imaging, there are several factors that need to be considered: (1)the size of the field of view; (2)the aberration correction of the system; (3)the selection of reconstruction parameters. As shown in Results section, the field of view (FOV) we used is × 25 *µm* × 25 *µm*, with the pixel size of 83.6 nm. If the FOV is expanded, the phase of 4Pi PSF will be field-dependent, due to the varying optical path differences from the edge to the center. This confined FOV results prevent from high-throughput imaging. By introducing vectorial PSF phase retrieval [30, 31] in separate regions, this problem may be improved. Additionally, fluorescence interference is highly sensitive to aberrations, which can lead to the deformation of the PSF and, consequently, distorted results with artifacts. Here we used adaptive optics, typically deformable mirrors [32], to address this problem. Although there was still a slight tilt in the PSF, it can have little influence on the quality of the reconstruction.

To overcome the non-uniformity of synthetic OTF in SIM reconstruction, which tends to cause artifacts and noises, a space-domain optimization was developed based on sparsity(*l*_1_ norm) and continuity (Frobenius Hessian norm). They can suppress the background and noise and meanwhile, slightly enhance isotropic resolution. By employing the ADMM algorithm for iteration, rapid convergence can be achieved.

## 4 Conclusion

In summary, we report I^2^SIM, a powerful approach to enhance axial resolution in imaging subcellular structures and morphologies of cells, which improves the axial resolution to ∼ 130 *nm*, about 2-fold improvement compared with 3D-SIM and maintains the lateral resolution of 3D-SIM at the same time. Using this approach, the fine axial structures that were originally difficult to discern in ordinary SIM can now be resolved, enhancing the optical sectioning capability. Despite the impressive axial resolution, it is not identical to the lateral resolution, and there is still room for improvement. By integrating innovative illumination pattern, such as four-beam interference, with advanced algorithms like deep learning, this issue can be addressed in the future.

## 5 Experimental Section

### Data reconstruction

To comply with the Nyquist smapling theorem, the axial scanning interval was set to 50 nm when collecting 3D Stacks of biological samples, given that the theoretical axial resolution is approximately 130 nm. Taking into account the thickness of the cells and the range of microtubules, the number of scanning layers was set to 21 and 31 respectively. This allows for reconstructing the raw data with minimal information loss while also minimizing photobleaching as much as possible.

The selection of reconstruction parameters plays a crucial role in the outcome of the reconstruction [29]. For the SIM reconstruction algorithm, we mainly adjusted the notch depth of *filter*_1_ and *filter*_2_, to surpass the background noise and maintain signals. In practical imaging process, we set the notch depth to 0.96 for *filter*_1_ and 0.98 for *filter*_2_.

For the optimization process, the parameters decide whether the optimization can converge and reach the best solution. λ_1_ determines the sparsity of the results. When the larger λ_1_, the noise is suppressed to a greater extent while non discontinuity is also enhanced with sharper edges. λ_2_ determines the significance of the Frobenius Hessian norm, and enhance the continuity of samples while sacrificing resolution. Correct configuration allows for ideal resolution and mitigated artifacts at the same time. The chose of parameters, however, depends on empirical trials, which is a drawback of the algorithm. For samples which are sparsely distributed in the FOV like fluorspheres, λ_1_ ought to be relatively large and for continuous samples like microtubules or actin filaments, a larger λ_2_ would be appropriate. λ_*z*_ controls the importance of axial continuity, which is set to 1 as default value. In biological imaging of this article, λ_1_ was 0.03, λ_2_ was 0.01, and λ_*z*_ was 1 to get a more continuous result.

### Sample preparation

25 mm diameter round precision glass coverslips (Thorlabs) were immersed in ultrapure water and vibrated in an ultrasonic cleaner for 15 minutes. Coverslips were sterilized using 95% ethanol. BSC-1 cells were purchased from the American Type Culture Collection and cultured in Minimum Essential Media (MEM; Thermo Fisher Scientific, Inc.) supplemented with 10% (v/v) fetal bovine serum (FBS; Thermo Fisher Scientific, Inc.). U2OS cells were purchased from the American Type Culture Collection and cultured in McCoy’s 5A Media (Thermo Fisher Scientific, Inc.) supplemented with 10% FBS. Cultures were maintained at 37 ^°^C in a humidified environment containing 5% CO_2_.

For bead samples, we used red fluorescent FluoSpheres beads (F8801) for fluorescent bead samples, which were maximally excited at 589 nm. The beads were diluted in ultrapure water at a ratio of 1:10, and vibrated in an ultrasonic cleaner for 5 min. Next, 200 *µ*l of the diluted beads were placed on a coverslip and allowed to stand for 10 min. The absorbent tissue was then used to remove excess water and Prolong Glass Antifade (Thermo Fisher Scientific, Inc.) before sealing the coverslip.

For the immunolabeling of microtubules, BSC-1 cells were seeded into the prepared coverslips. After overnight incubation, the cells were washed thrice with phosphate-buffered saline (PBS, Thermo Fisher Scientific, Inc.), fixed with 3% (m/v) paraformaldehyde (PFA, Electron Microscopy Sciences) and 0.1% (v/v) glutaraldehyde (GA, SigmaAldrich Co., LLC) for 13 minutes at 37 ^°^C, and quenched with NaHB_4_ for 7 minutes at room temperature (RT). The cells were incubated with 0.2% (v/v) Triton X-100 (Sigma-Aldrich Co., LLC) and 5% goat serum (Thermo Fisher Scientific, Inc.) for 1 hour at RT. Microtubules were stained with mouse alpha and beta tubulin antibodies (WA31679510 and TG2597441, Invitrogen) at 1:1:200 dilution in PBS overnight at 4 ^°^C and goat anti-mouse STAR Red (Abberior GmbH) at 1:200 dilution in PBS for 1 hour. Samples were washed three times with PBS before imaging, with two washes for 3 minutes and the last wash for 15 minutes.

To label the actin filaments in U2OS cells, the cells were seeded onto the prepared coverslips. After overnight incubation, the cells were washed thrice with PBS. Next, the cells were fixed with 3% PFA and 0.1% GA in PBS for 13 minutes and then rinsed three times with PBS. Subsequently, the cells were incubated with iFluor 555 phalloidin (Yeasen Biotechnology Co., Ltd.) diluted at a concentration of 1.5:200 for 30 minutes at room temperature. The cells were then washed three times with PBS with two washes for 3 minutes and the last wash for 15 minutes and stored in PBS at 4 ^°^C until imaged.

To label the mitochondrial outer membrane in U2OS cells, the cells were seeded onto the prepared coverslips. After overnight incubation, the cells were washed once with PBS, then fixed with 3% (m/v) PFA and 0.1% (v/v) GA for 13 minutes at 37 ^°^C, and quenched with NaHB_4_ for 7 minutes at RT. Subsequently, The cells were incubated with 0.2% (v/v) Triton X-100 (Sigma-Aldrich Co., LLC) and 5% goat serum (Thermo Fisher Scientific, Inc.) for 50 minutes at RT. The cells were then incubated overnight at 4 ^°^C with rabbit anti-TOMM20 (Thermo Fisher Scientific, 1:100). Following three washes for 3 minutes each with wash buffer, the cells were incubated with Alexa Fluor 546 (AF546) (Thermo Fisher Scientific, 0.5:250) for 1 hour at RT. The cells were then washed three times with PBS with two washes for 3 minutes and the last wash for 15 minutes and stored in PBS at 4 ^°^C until imaged.

## Supporting Information

Supporting Information is available from the Wiley Online Library or from the author.

The code for isotropic Frobenius Hessian optimization in I^2^SIM and one raw SIM data are available at https://github.com/ZJUOPTKuangLab/I2SIM.

## Acknowledgements

This work was financially sponsored by the National Natural Science Foundation of China (62125504, 62361166631); STI 2030—Major Projects (2021ZD0200401); the Fundamental Research Funds for the Central Universities (2022FZZX01-20).

## Conflict of Interest

The authors declare no conflict of interest.

## Notes

### Competing Interest Statement

The authors have declared no competing interest.

### Summary of Updates

Results of the mitochondrial outer membrane have been added.

## References

[1] M.G.L. Gustafsson, Journal of Microscopy 2000, 198, 2 82.

[2] M. G. Gustafsson, L. Shao, P. M. Carlton, C. J. R. Wang, I. N. Golubovskaya, W. Z. Cande, D. A. Agard, J. W. Sedat, Biophysical Journal 2008, 94, 12 4957.

[3] L. Shao, P. Kner, E. H. Rego, M. G. L. Gustafsson, Nature Methods 2011, 8, 12 1044.

[4] X. Chen, S. Zhong, Y. Hou, R. Cao, W. Wang, D. Li, Q. Dai, D. Kim, P. Xi, Light: Science & Applications 2023, 12, 1 172.

[5] H. Zhu, Y. Sun, L. Yin, J. Han, M. Cai, Q. Liu, C. Kuang, X. Hao, X. Liu, Optics Express 2021, 29, 14 21428.

[6] M. Cai, H. Zhu, Y. Sun, L. Yin, F. Xu, H. Wu, X. Hao, R. Zhou, C. Kuang, X. Liu, Optics Express 2022, 30, 5 7938.

[7] G. Wen, S. Li, Y. Liang, L. Wang, J. Zhang, X. Chen, X. Jin, C. Chen, Y. Tang, H. Li, PhotoniX 2023, 4, 1 19.

[8] G. Wen, S. Li, L. Wang, X. Chen, Z. Sun, Y. Liang, X. Jin, Y. Xing, Y. Jiu, Y. Tang, H. Li, Light: Science & Applications 2021, 10, 1 70.

[9] J. Qian, K. Xu, S. Feng, Y. Liu, H. Ma, Q. Chen, C. Zuo, ACS Photonics 2024.

[10] R. Cao, Y. Chen, W. Liu, D. Zhu, C. Kuang, Y. Xu, X. Liu, Biomedical Optics Express 2018, 9, 10 5037.

[11] K. Wicker, O. Mandula, G. Best, R. Fiolka, R. Heintzmann, Optics Express 2013, 21, 2 2032.

[12] J. Qian, Y. Cao, Y. Bi, H. Wu, Y. Liu, Q. Chen, C. Zuo, eLight 2023, 3, 1 4.

[13] Z. Wang, T. Zhao, H. Hao, Y. Cai, K. Feng, X. Yun, Y. Liang, S. Wang, Y. Sun, P. R. Bianco, K. Oh, M. Lei, Advanced Photonics 2022, 4, 2 026003.

[14] M. P. Strauss, A. T. F. Liew, L. Turnbull, C. B. Whitchurch, L. G. Monahan, E. J. Harry, PLOS Biology 2012911, 10, 9 e1001389.

[15] V. W. Rowlett, W. Margolin, Biophysical Journal 2014, 107, 8 L17.

[16] V. C. Cogger, G. P. McNerney, T. Nyunt, L. D. DeLeve, P. McCourt, B. Smedsrød, D. G. Le Couteur, T. R. Huser, Journal of Structural Biology 2010, 171, 3 382.

[17] L. Rodermund, H. Coker, R. Oldenkamp, G. Wei, J. Bowness, B. Rajkumar, T. Nesterova, D. M. Susano Pinto, L. Schermelleh, N. Brockdorff, Science 2021, 372, 6547 eabe7500.

[18] X. Li, Y. Wu, Y. Su, I. Rey-Suarez, C. Matthaeus, T. B. Updegrove, Z. Wei, L. Zhang, H. Sasaki, Y. Li, M. Guo, J. P. Giannini, H. D. Vishwasrao, J. Chen, S.-J. J. Lee, L. Shao, H. Liu, K. S. Ramamurthi, J. W. Taraska, A. Upadhyaya, P. La Riviere, H. Shroff, Nature Biotechnology 2023, 41, 9 1307.

[19] L. Shao, B. Isaac, S. Uzawa, D. A. Agard, J. W. Sedat, M. G. L. Gustafsson, Biophysical Journal 2008, 94, 12 4971.

[20] L. Shao, L. Winoto, D. Agard, M. Gustafsson, J. Sedat, Journal of Microscopy 2012, 246, 3 229.

[21] Z. Ouyang, Q. Wang, X. Li, Q. Dai, M. Tang, L. Shao, W. Gou, Z. Yu, Y. Chen, B. Zheng, L. Chen, C. Ping, X. Bi, B. Xiao, X. Yu, C. Liu, L. Chen, J. Fan, X. Huang, Y. Zhang, Nature Methods 2024, 1–13.

[22] Gustafsson, Agard, Sedat, Journal of Microscopy 1999, 195, 1 10.

[23] Y. Sun, H. Zhu, L. Yin, H. Wu, M. Cai, W. Sun, Y. Xu, X. Yang, J. Han, W. Liu, Y. Han, X. Hao, R. Zhou, C. Kuang, X. Liu, Advanced Photonics 2023, 5, 5 056007.

[24] Y. Sun, H. Zhu, X. Yang, E. He, H. Wu, L. Yin, W. Sun, X. Luo, Y. Han, X. Hao, R. Zhou, C. Kuang, X. Liu, ACS Photonics 2025, 12, 1 419.

[25] X. Huang, J. Fan, L. Li, H. Liu, R. Wu, Y. Wu, L. Wei, H. Mao, A. Lal, P. Xi, L. Tang, Y. Zhang, Y. Liu, S. Tan, L. Chen, Nature Biotechnology 2018, 36, 5 451.

[26] W. Zhao, S. Zhao, L. Li, X. Huang, S. Xing, Y. Zhang, G. Qiu, Z. Han, Y. Shang, D.-e. Sun, C. Shan, R. Wu, L. Gu, S. Zhang, R. Chen, J. Xiao, Y. Mo, J. Wang, W. Ji, X. Chen, B. Ding, Y. Liu, H. Mao, B.-L. Song, J. Tan, J. Liu, H. Li, L. Chen, Nature Biotechnology 2022, 40, 4 606.

[27] R. Cao, Y. Li, X. Chen, X. Ge, M. Li, M. Guan, Y. Hou, Y. Fu, X. Xu, C. Leterrier, S. Jiang, B. Gao, P. Xi, Nature Methods 2023, 20, 8 1183.

[28] S. Lefkimmiatis, A. Bourquard, M. Unser, IEEE Transactions on Image Processing 2012, 21, 3 983.

[29] S. Boyd, Foundations and Trends® in Machine Learning 2010, 3, 1 1.

[30] X. Yang, H. Zhu, Y. Sun, H. Wu, Y. Han, X. Hao, C. Kuang, X. Liu, Photonics Research 2024, 12, 11 2447.

[31] N. H. Thao, O. Soloviev, M. Verhaegen, JOSA A 2020, 37, 1 16.

[32] F. Huang, G. Sirinakis, E. S. Allgeyer, L. K. Schroeder, W. C. Duim, E. B. Kromann, T. Phan, F. E. Rivera-Molina, J. R. Myers, I. Irnov, M. Lessard, Y. Zhang, M. A. Handel, C. Jacobs-Wagner, C. P. Lusk, J. E. Rothman, D. Toomre, M. J. Booth, J. Bewersdorf, Cell 2016, 166, 4 1028.

