## Supporting Information for "I^2^SIM: Boosting High-Fidelity Isotropic Super-Resolution with Image Interference and Spatial-Spectral Optimization"

#### 1 Details of optimization algorithm

We adopted ADMM to solve the optimization problem in **Equation 5**. Scaled ADMM was employed for simplicity. The Augmented Lagrangian function was constructed as:

$$\begin{aligned} L =& \frac{1}{2}\|Hf-b\|_2^2 + \lambda_1\|d_1\|_1 + \\ & \lambda_2\|\sqrt{d_{xx}^2+d_{xy}^2+d_{yy}^2+d_{xz}^2+d_{yz}^2+d_{zz}^2}\|_1 + \\ & \sum_{\text{for all } d}\left[\frac{\rho}{2}\|Af-d+\omega\|_2^2 - \frac{\rho}{2}\|\omega\|_2^2\right]. \end{aligned} \tag{1}$$

For each $d(d_1,\ d_{xx},\ d_{xy},\ d_{yy},\ d_{xz},\ d_{yz},\ d_{zz})$, there is one $A$ operator and one scaled dual variable $\omega$. $A$ is in the form of $I,\ \nabla_{xx},\ \sqrt{2}\nabla_{xy},\ \nabla_{yy},\ \sqrt{2\lambda_z}\nabla_{xz},\ \sqrt{2\lambda_z}\nabla_{yz},\ \lambda_z\nabla_{zz}$, respectively. Here, $I$ denotes an identity matrix of the same size as the input image stack, and operator $\nabla$ represents the second-order partial derivative operation on the image stack indicated by its subscripts.

By minimizing $L$ with respect to $f$, $d$, and $\omega$ iteratively, the best value of $f$ can be found. Take the partial derivative of $L$ with respect to $f$ and set it equal to zero, we can get:

$$\frac{\partial L}{\partial f} = H^T(Hf-b) + \sum_{\text{for all } d}\left[\rho A^T(Af-d+\omega)\right] = 0. \tag{2}$$

$\cdot^T$ denotes the transpose of a matrix. $f$ can be extracted and solved using the Fourier Transform in **Equation S3**.

$$f^{i+1} = ifftn\left[\frac{\sum_{\text{for all } d}\tilde{A}^*\odot(\tilde{d}^i-\tilde{\omega}^i)+\frac{\tilde{H}^*\odot\tilde{b}}{\rho}}{\sum_{\text{for all } d}|\tilde{A}|^2+\frac{|\tilde{H}|^2}{\rho}}\right], \tag{3}$$

$\tilde{\cdot}$ is the Fourier Transform of matrix, $\cdot^*$ represents the conjugate of complex matrix, and $\odot$ denotes element-wise multiplication.

Next we update each $d$ through the same way as $f$, for all $d$ except $d_1$, the partial derivative of $L$ is calculated(**Equation S4)**).

$$\frac{\partial L}{\partial d} = \frac{\lambda_2 d}{\sqrt{d_{xx}^2+d_{xy}^2+d_{yy}^2+d_{xz}^2+d_{yz}^2+d_{zz}^2}} + \rho(d-C^i) = 0 \tag{4}$$

$$\begin{cases} C_{xx}^i = \nabla_{xx}f^{i+1}+\omega_{xx}^i \\ C_{xy}^i = \sqrt{2}\nabla_{xy}f^{i+1}+\omega_{xy}^i \\ C_{yy}^i = \nabla_{yy}f^{i+1}+\omega_{yy}^i \\ C_{xz}^i = \sqrt{2\lambda_z}\nabla_{xz}f^{i+1}+\omega_{xz}^i \\ C_{yz}^i = \sqrt{2\lambda_z}\nabla_{yz}f^{i+1}+\omega_{yz}^i \\ C_{zz}^i = \lambda_z\nabla_{zz}f^{i+1}+\omega_{zz}^i \end{cases}$$

Since the $d$'s are coupled, we need to find the multivariate function extremum of $L$ with respect to $d$. When the six equations derived from Equation S3 are combined, the following relationships are obtained:

$$\frac{d_{xx}}{C_{xx}} = \frac{d_{xy}}{C_{xy}} = \frac{d_{yy}}{C_{yy}} = \frac{d_{xz}}{C_{xz}} = \frac{d_{yz}}{C_{yz}} = \frac{d_{zz}}{C_{zz}}. \tag{5}$$

Take $d_{xx}$ as example, substituting the above conclusions(**Equation S5**) back into Equation S4, we can obtain:

$$\frac{\lambda_2 d_{xx}}{\sqrt{\sum_{\text{for all } C}\left(\frac{d_{xx}}{C_{xx}}C\right)^2}} + \rho(d_{xx} - C_{xx}) = 0, \tag{6}$$

which can be simplified to:

$$\frac{\lambda_2 \cdot sign(d_{xx})|C_{xx}|}{\sqrt{\sum_{\text{for all } C}(C)^2}} + \rho(d_{xx} - C_{xx}) = 0. \tag{7}$$

Here, $sign(d)$ is 1 when $d > 0$, -1 when $d < 0$, and 0 when $d = 0$. Through classification discussion, the iterative formula for $d_{xx}$ can be summarized as a soft-threshold function:

$$d_{xx}^{i+1} = \begin{cases} C_{xx}^i - |C_{xx}^i|\frac{\lambda_2/\rho}{\sqrt{\sum(C)^2}}, & C_{xx}^i > |C_{xx}^i|\frac{\lambda_2/\rho}{\sqrt{\sum(C)^2}} \\ 0, & -|C_{xx}^i|\frac{\lambda_2/\rho}{\sqrt{\sum(C)^2}} < C_{xx}^i < |C_{xx}^i|\frac{\lambda_2/\rho}{\sqrt{\sum(C)^2}} \\ C_{xx}^i + |C_{xx}^i|\frac{\lambda_2/\rho}{\sqrt{\sum(C)^2}}, & C_{xx}^i < -|C_{xx}^i|\frac{\lambda_2/\rho}{\sqrt{\sum(C)^2}} \end{cases} \tag{8}$$

Similar operations can be applied to the iteration of other $d$'s (except $d_1$). By considering case when $d = 0$, Eq. **??** can be transformed into a more general form:

$$d_{xx}^{i+1} = max\left(\sqrt{\sum(C^i)^2} - \frac{\lambda_2}{\rho}\right) \odot \frac{C_{xx}^i}{\sqrt{\sum(C^i)^2}}. \tag{9}$$

For $d_1$, it is easy to derive the iteration formula of $d_1$, which is also a soft-threshold function:

$$d_1^{i+1} = S_{\frac{\lambda_1}{\rho}}(f^{i+1} + \omega^i), \tag{10}$$

where $S_\gamma(y) = max(y - \gamma, 0) + min(y + \gamma, 0)$. For all $\omega$, the iterations are similar, which can be denoted as:

$$w^{i+1} = w^i + Af^{i+1} - d^{i+1} \tag{11}$$

The pseudo code is as follows:

**Algorithm 1** Frobenius Hessian Sparsity Optimization

**Input:** $f^0,\ H,\ \lambda_1,\ \lambda_2,\ \lambda_z,\ \rho$, IterNum
**Output:** de-noised image stack $f^*$ with depressed artifacts

```
1: Initialize:
2: all d, ω ← f^0.
3: for i = 0 to IterNum-1 do
4:     Update f^{i+1} according to Equation S3.
5:     Update d_1^{i+1} according to Equation S10.
6:     Update d_xx^{i+1}–d_zz^{i+1} according to Equation S9.
7:     Update all ω^{i+1} according to Equation S11.
8: end for
9: f* ← f^{IterNum}
```

### 2 Details in SIM reconstruction

#### 2.1 ordinary Wiener-SIM

3D-SIM uses three beams to illuminate samples, generating a three dimension pattern, which can be expressed as:

$$I_{\theta,\varphi}(\vec{r},\vec{z}) = I_0[1 + 2m^2 + 4m \cdot cos(2\pi\vec{p_z} \cdot \vec{z}) \cdot cos(2\pi\vec{p_{x,y}} \cdot \vec{r} + \varphi) + 2m^2 cos(4\pi\vec{p_{x,y}} \cdot \vec{r} + 2\varphi)] \tag{12}$$

Here, $I_0$ denotes the intensity of illumination, and $m$ denotes the modulation depth of the pattern in the focal plane. $p_{x,y} = \frac{nsin\phi}{\lambda}$, $p_z = \frac{n(1-cos\phi)}{\lambda}$, are the spatial frequency of pattern in the lateral and axial plane, and $\phi = arcsin(NA/n)$ is the polar angle of incidence, NA is the numerical aperture, $\lambda$ is the excitation wavelength, n is the refractive index of the immersion medium. $\vec{r} = cos\theta\vec{x} + sin\theta\vec{y}$ denotes the lateral position vector. $\theta$ denotes the orientation of the illumination pattern's vector in the lateral plane. $\varphi$ denotes the phase of the illumination pattern. The illumination pattern has three harmonics with the relative weight of $a_0 = 1 + 2m^2$, $a_1 = 4m$, $a_2 = 2m^2$.
We use $S(\vec{r}, \vec{z})$ to denote the distribution of the fluorophores' emission light distribution, and the image stack acquired by sCMOS of one orientation $\theta$ and one phase $\varphi$ can be expressed as:

$$D_{\theta,\varphi}(\vec{r}, \vec{z}) = [S(\vec{r}, \vec{z} \cdot I_{\theta,\varphi}(\vec{r}, \vec{z})] \otimes H(\vec{r}, \vec{z}) \tag{13}$$

$H(\vec{r}, \vec{z})$ is the 3D point spread function (PSF). Then, we apply 3D Fourier Transform on both side. Because the axial scanning is achieved by moving the sample, the axial frequency of the illumination pattern and the PSF are coupled together. The 3D frequency domain expression can be given by:

$$\begin{aligned} D_{\theta,\varphi}(\vec{k_r}, \vec{k_z}) =& [S(\vec{k_r}, \vec{k_z} \otimes I_{\theta,\varphi}(\vec{k_r}, \vec{k_z})] \cdot H(\vec{k_r}, \vec{k_z}) \\ =& I_0 \left\{ a_0 S(\vec{k_r}, \vec{k_z}) + a_2 \left[ S(\vec{k_r} - 2\vec{p_r}, \vec{k_z}) e^{j2\varphi} + S(\vec{k_r} + 2\vec{p_r}, \vec{k_z}) e^{-j2\varphi} \right] \right\} H(\vec{k_r}, \vec{k_z}) + \\ & a_1 \left[ S(\vec{k_r} - \vec{p_r}, \vec{k_z}) e^{j\varphi} + S(\vec{k_r} + \vec{p_r}, \vec{k_z}) e^{-j\varphi} \right] \cdot \left[ H(\vec{k_r}, \vec{k_z} - \vec{p_z}) + H(\vec{k_r}, \vec{k_z} + \vec{p_z}) \right] \end{aligned} \tag{14}$$

There are five unknown frequency components in **Equation S14**, and five different $\varphi$ are needed to resolve them. Then we get five equations in the format of the matrix:

$$\begin{bmatrix} D_{\theta,\varphi_1}(\vec{k}) \\ D_{\theta,\varphi_2}(\vec{k}) \\ D_{\theta,\varphi_3}(\vec{k}) \\ D_{\theta,\varphi_4}(\vec{k}) \\ D_{\theta,\varphi_5}(\vec{k}) \end{bmatrix} = I_0 A \begin{bmatrix} S(\vec{k_r}, \vec{k_z}) \cdot H(\vec{k_r}, \vec{k_z}) \\ S(\vec{k_r} - \vec{p_r}, \vec{k_z}) \cdot [H(\vec{k_r}, \vec{k_z} - \vec{p_z}) + H(\vec{k_r}, \vec{k_z} + \vec{p_z})] \\ S(\vec{k_r} + \vec{p_r}, \vec{k_z}) \cdot [H(\vec{k_r}, \vec{k_z} - \vec{p_z}) + H(\vec{k_r}, \vec{k_z} + \vec{p_z})] \\ S(\vec{k_r} - 2\vec{p_r}, \vec{k_z}) \cdot H(\vec{k_r}, \vec{k_z}) \\ S(\vec{k_r} + 2\vec{p_r}, \vec{k_z}) \cdot H(\vec{k_r}, \vec{k_z}) \end{bmatrix} = I_0 A \begin{bmatrix} C_0(\vec{k}) \\ C_{-1}(\vec{k}) \\ C_{+1}(\vec{k}) \\ C_{-2}(\vec{k}) \\ C_{+2}(\vec{k}) \end{bmatrix} \tag{15}$$

Here, $A$ is the coefficient matrix and can be expressed as:

$$A = \begin{bmatrix} a_o & a_1 e^{j\varphi_1} & a_1 e^{-j\varphi_1} & a_2 e^{j2\varphi_1} & a_2 e^{-j2\varphi_1} \\ a_o & a_1 e^{j\varphi_2} & a_1 e^{-j\varphi_2} & a_2 e^{j2\varphi_2} & a_2 e^{-j2\varphi_2} \\ a_o & a_1 e^{j\varphi_3} & a_1 e^{-j\varphi_3} & a_2 e^{j2\varphi_3} & a_2 e^{-j2\varphi_3} \\ a_o & a_1 e^{j\varphi_4} & a_1 e^{-j\varphi_4} & a_2 e^{j2\varphi_4} & a_2 e^{-j2\varphi_4} \\ a_o & a_1 e^{j\varphi_5} & a_1 e^{-j\varphi_5} & a_2 e^{j2\varphi_5} & a_2 e^{-j2\varphi_5} \end{bmatrix} \tag{16}$$

The five frequency components on the right side of Equation 15 can then be calculated by:

$$\begin{bmatrix} C_0(\vec{k}) \\ C_{-1}(\vec{k}) \\ C_{+1}(\vec{k}) \\ C_{-2}(\vec{k}) \\ C_{+2}(\vec{k}) \end{bmatrix} = \frac{1}{I_0} A^{-1} \begin{bmatrix} D_{\theta,\varphi_1}(\vec{k}) \\ D_{\theta,\varphi_2}(\vec{k}) \\ D_{\theta,\varphi_3}(\vec{k}) \\ D_{\theta,\varphi_4}(\vec{k}) \\ D_{\theta,\varphi_5}(\vec{k}) \end{bmatrix} \tag{17}$$

Here $(\cdot)^{-1}$ denotes the operation of the matrix inverse transform. Performing the same operation for the other two orientations $\theta_2$ and $\theta_3$, we can get 15 separated frequency components. The final super-resolution image can be obtained using Wiener filter:

$$S_{SR}(\vec{k}) = \frac{\sum_{\theta,l} H^*_{l,\theta}(\vec{k}) C_{l,\theta}(\vec{k_r} + l \cdot \vec{p}_{r,\theta}, \vec{k_z})}{\sum_{\theta,l} |H_{l,\theta}(\vec{k})|^2 + w^2} A(\vec{k}) \tag{18}$$

Where $S_{SR}(\vec{k}$ is the super-resolution spectrum. $l$ (-2, -1, 0, 1, 2) denotes the order of the separated frequency components, and $H_{l,\theta}(\vec{k})$ is the corresponding OTF which is shifted to the $l$-th place in the frequency domain of orientation $\theta$. For each orientation $\theta$, when $l = 0, \pm 2$, $H_{l,\theta}(\vec{k}) = H(\vec{k_r} + l \cdot \vec{p}_{r,\theta}, \vec{k_z})$, when $l = \pm 1$, $H_{l,\theta}(\vec{k}) = H(\vec{k_r} + l \cdot \vec{p}_{r,\theta}, \vec{k_z} - \vec{p_z}) + H(\vec{k_r} + l \cdot \vec{p}_{r,\theta}, \vec{k_z} + \vec{p_z})$. $(\cdot)^*$ is the symbol of complex conjugate. $w^2$ is the Wiener parameter which was taken to be a constant and adjusted empirically.

#### 2.2 Preliminary spectral processing using notch filter

Wiener-SIM will cause the synthetic spectrum abnormal, resulting in sidelobe artifacts. What's more, peaks at the center of each order frequency component can amplify the background noise. So we adapt spectral modulation to solve this problem. Before we apply two spectrum filters on the synthetic spectrum as mentioned in the main text, a notch filter is used to suppress the high-frequency peak. This will constitute the first step of spectrum optimization.

$$C_{SR} = \sum_{\theta,l} C_{l,\theta}(\vec{k}_r + l \cdot \vec{p}_{r,\theta}, \vec{k}_z) \cdot notch_{l,\theta}(\vec{k}) \cdot H^{att}_{l,\theta}(\vec{k}) \tag{19}$$

Where $notch_{0,\theta}(\vec{k}) = 1 - d \cdot exp\left[\left(\frac{\vec{k}_x^2+\vec{k}_y^2}{|\vec{p}_r|^2} + \frac{\vec{k}_z^2}{|\vec{p}_z|^2}\right)/2/width\right]$. For $l = \pm 1$, $notch_{l,\theta}(\vec{k}) = notch_{0,\theta}(\vec{k}_r + l \cdot \vec{p}_{r,\theta}, \vec{k}_z + \vec{p}_z) + notch_{0,\theta}(\vec{k}_r + l \cdot \vec{p}_{r,\theta}, \vec{k}_z - \vec{p}_z)$, for $l = \pm 2$, $notch_{l,\theta}(\vec{k}) = notch_{0,\theta}(\vec{k}_r + l \cdot \vec{p}_{r,\theta}, \vec{k}_z)$. $att$ is the frequency attenuation, and $d$ and $width$ are the notch depth and width, respectively. Then the $C_{SR}$ can be used for further processing.

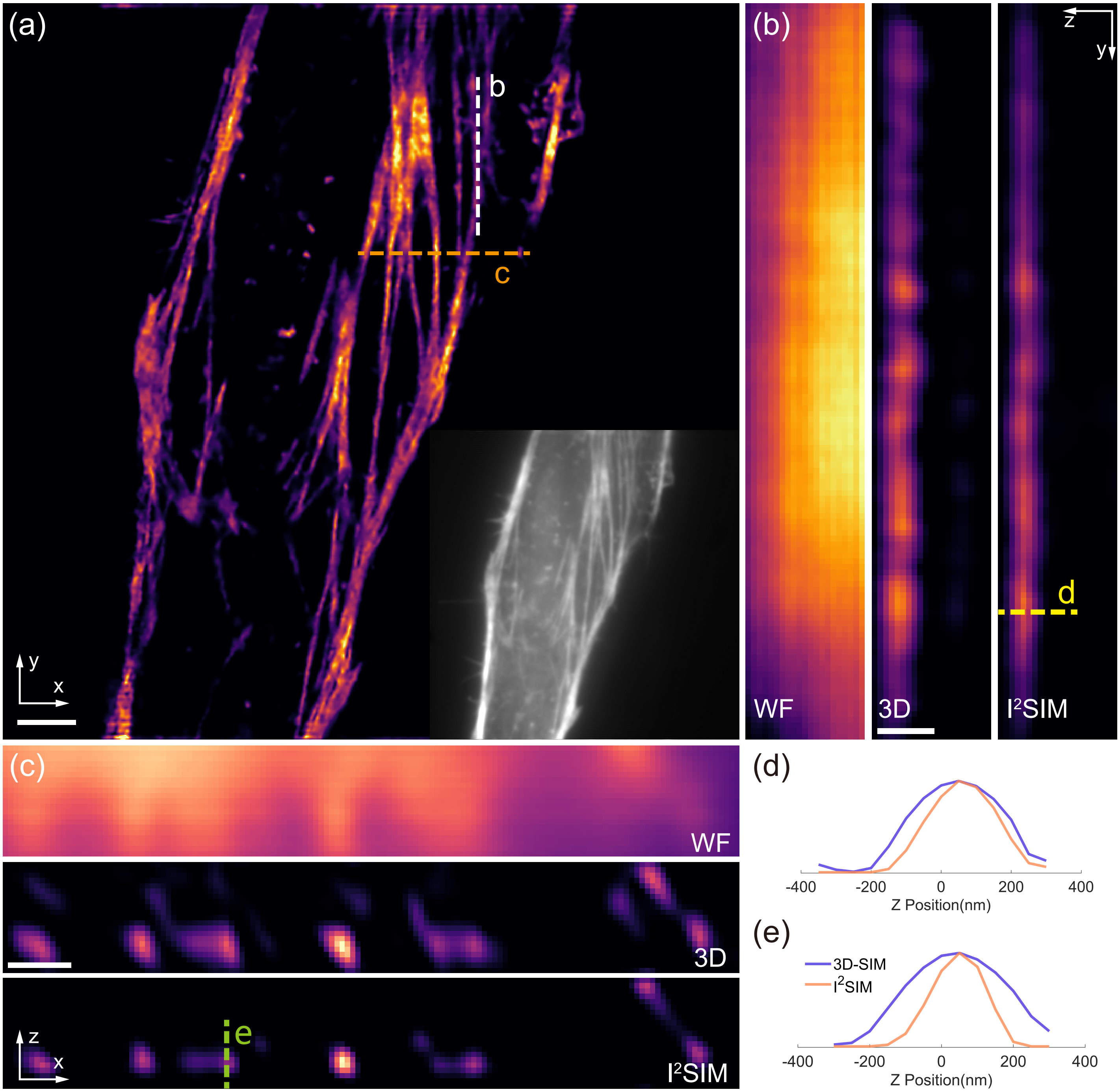


Figure S1: $I^2$SIM enables the enhancement of axial resolution for actin filaments. a) Single lateral plane of actin filaments in U2OS cells labeled with iFluor 555 phalloidin. b) Axial views corresponding to the white dashed line in (a), comparing wide-field microscopy(WF), 3D-SIM(3D), $I^2$SIM. c) Axial views corresponding to the orange dashed line in (a), comparing wide-field microscopy(WF), 3D-SIM(3D), $I^2$SIM. d) Line profiles corresponding to the yellow dashed line in (b). e) Line profiles corresponding to the green dashed line in (c). Scale bars, 2 $\mu$m (a); 500 nm (b,c).
